# Angiogenesis is critical for the regenerative effects of exercise

**DOI:** 10.1101/2021.11.17.468965

**Authors:** Supriya S. Wariyar, Alden D. Brown, Tina Tian, Tana S Pottorf, Patricia J. Ward

## Abstract

Enhancing axon regeneration is a major focus of nerve injury research, and the quality of the surgical nerve repair plays a large role in the aggregate success of nerve regeneration. Additionally, exercise is known to promote successful axon regeneration after surgical nerve repair. In this study, we asked how exercise-induced nerve regeneration is affected when a transected nerve is repaired with or without fibrin glue. Fibrin glue repaired nerves exhibited greater vasculature within the tissue bridge compared to nerves that were intrinsically repaired. Fibrin glue repaired nerves also exhibited more robust axon regeneration after exercise compared to nerves that were not repaired with fibrin glue. When angiogenesis of the tissue bridge was prevented, exercise was unable to enhance regeneration despite the presence of fibrin glue. These findings suggest that the biological properties of fibrin glue enhance angiogenesis within the repair site, and a vascularized bridge is required for enhanced axon elongation with exercise. The combination of fibrin glue repair and exercise resulted in notable differences in vascular growth, axon elongation, neuromuscular junction reinnervation, and functional recovery. Fibrin glue should be considered as an adjuvant for nerve repair to enhance the subsequent efficacy of activity- and physical therapy-based treatment interventions.

## Introduction

After injury, peripheral nerves exhibit a limited capacity for regeneration, and the success of axon regeneration relies on the temporal coordination of both resident cells (injured neurons and repair Schwann cells) and cells that are recruited to the injury site (macrophages, fibroblasts, epithelial cells, and Schwann cells) (1, 2). During regeneration, Schwann cells lead the axons across an ill-defined “bridge” that forms in the gap between the two nerve stumps after a transection (3, 4). The construction of this tissue bridge is initiated by hypoxia-sensitive macrophages that attract endothelial cells from the distal and proximal nerve stumps in a VEGFA-dependent mechanism (5). These endothelial cells re-vascularize the tissue bridge and provide a physical substrate for the migration of Schwann cells and the formation of cords, known as Bands of Büngner, which then attract the regenerating axons. Some axons will “wander, resulting in axonal escape or misdirection (6, 7), but axons that follow the Schwann cell cords across the tissue bridge can successfully reenter an endoneurial tube destined for a peripheral target (8, 9). Importantly, if the nascent vessels are redirected, so too are the Schwann cells and axons, indicating the importance of the physical pathway provided by re-vascularization of the tissue bridge (5).

Although peripheral nerves have this limited potential to regenerate, functional recovery is quite challenging in many clinical scenarios, such as when the injury is closer to the spinal cord, is far away from the target organ, involves mixed (motor and sensory) nerves, and/or involves gaps ≥ 5 mm. The reason for poor functional recovery is generally attributed to the inability of injured neurons to regenerate their axons over extremely long distances at the very slow rate of ∼1 mm/day (2, 10). Treatment approaches are surgical, and the gold standard is to utilize microsutures to hold the cut nerve stumps in close proximity. Microsutures must be applied with great care because they can be associated with inflammation, fascicle ligation by the suture, and neuroma formation around the sutures (10, 11). Alternatively, fibrin glue has been used to form a clot around the nerve stumps, demonstrating equivalent or superior results compared to suture alone (12). However, the use of fibrin glue clinically is questionable due to concerns regarding potential immunogenicity, dehiscence due to poor tensile strength when used alone, and lack of FDA approval for the indication of nerve repair (12–14). Each nerve repair is unique to the style and preference of the operating surgeon, as well as the nature and location of the nerve injury. While microsutures are the gold standard, fibrin glue can be useful for small caliber nerves or when operating within small spaces, e.g., pediatric nerve repair.

Commercially-available fibrin sealants were developed in the 1970s, and the initial goals of the fibrin “suture,” or clot, were biomechanical stabilization of the approximated nerve ends and to provide the opportunity to hold fine adjustments of fascicles in place (15, 16). The process of clotting occurs naturally after disruption of the blood-nerve-barrier, and deposition of fibrin occurs within the CNS and PNS after spinal cord or nerve injury, respectively (17, 18). And while fibrin occurs naturally in clots, commercial fibrin glues are produced by the combination of various xenogenic factors, such as bovine and/or pooled human plasma, which have the potential for immunogenicity that might negatively affect regeneration by altering the immune response at the site of injury (12, 19).

Fibrin also has other known biological effects. For example, fibrin inhibits neurite growth *in vitro* and inhibits Schwann cell migration dose-dependently *in vivo* (17, 20). Fibrin also affects endothelial cell migration and the ability to form capillary-like structures (21–23), and as described above, angiogenesis by epithelial cells is a necessary contingent step for the development of Schwann cell cords (5). We and others have shown that running exercise – an activity-based experimental treatment – robustly enhances axon regeneration (24–27), and the mechanisms involved in this enhancement rely on both central and peripheral mechanisms, including angiogenesis, immunomodulation, and axon elongation. We hypothesized that the biological effects of fibrin glue may inhibit regeneration and reduce the benefits of exercise on axon regeneration and recovery. Thus, in the current study, we determined the effect of running exercise with or without fibrin glue repair on macrophage infiltration, blood vessel formation, axon elongation, and functional recovery after nerve transection.

## Materials and Methods

### Experimental Design

All experiments were conducted on adult (2-4 months old) mice weighing 18-30 grams using approximately equal males and females for each assay. All experiments were approved by the Institutional Animal Care and Use Committee of Emory University. *Thy-1-YFP-H* transgenic (Jackson laboratory stock no. 003782) or wildvtype mice *C57BL/6J* (Jackson laboratory stock no. 000664) were used. *Thy-1-YFP-H* mice express yellow fluorescent protein (YFP) at high levels in a subset of sensory and motor neurons. It was assumed that these YFP+ axons formed a representative sample of all axons in the nerve. A total of 19 nerves were studied from 11 wild type mice for immunohistochemistry of PECAM-1, IBA-1, and CD68 at 7 days post-injury. For the cabozantinib experiments (VEGF-R inhibitor), 8 nerves from 8 wild type mice were used to quantify vascularization of the injury site and axon regeneration at 7 days post-injury. To measure axon regeneration, a total of 29 nerves from 15 Thy-1-YFP-H mice were used to quantify axon profile lengths at 14-days post-injury. A total of 32 wild type mice were used for EMG studies at the 28-day time point. The gastrocnemius muscles from an additional 17 wild type mice were analyzed for NMJ reinnervation at the 28-day time point. The 7- and 14-day time points were chosen for most analyses as macrophage infiltration, angiogenesis within the tissue bridge, and axon crossing of the repair site are all acute events, and it is unlikely to detect histological differences at chronic timepoints (5, 28, 29). Functional recovery and reinnervation of muscle were performed at 28-days post-injury as earlier time points are unlikely to detect functional differences after this type of nerve injury and repair model.

### Surgical Procedure for Nerve Repair

Mice were assigned to untreated and exercised groups, and nerves were transected and repaired either with or without fibrin glue. During transection surgery, mice were subject to 5% isoflurane in 1L/min oxygen to inducing anesthesia and maintained at 2% isoflurane in 1L/min oxygen. The sciatic nerve was exposed by a minimal incision and blunt dissection through the hamstring fascia. The whole sciatic nerve was transected with sharp microscissors, making sure the proximal and distal stumps were in proximity and without disturbing surrounding fascia or muscle. In this manner, the fascia held the nerve stumps in their original orientation. We termed this an “intrinsic repair” referring to the natural formation of a tissue bridge followed by the regeneration of axons, as previously characterized (5). For fibrin glue repair, thrombin (MP Biomed, Catalog no. 154163) and fibrinogen (Sigma, Catalog no. F4129) in the ratio of 2:1, respectively, were added to the nerve stumps (30, 31). Muscle and skin were closed with suture. For acute analyses, surgeries were performed bilaterally. For functional recovery assays, surgeries were performed unilaterally. Animals survived 7, 14, or 28 days after surgery.

### Treadmill training

Beginning the third day post-nerve transection and repair, animals assigned to exercise groups were treadmill trained for five days per week using either an interval or continuous treadmill exercise paradigm that is effective in females and males, respectively (25, 26, 32, 33). Briefly, male mice were trained for one hour at a speed of 16.7 cm/sec, and female mice were trained for four two-minute repetitions at 33.33 cm/sec with a five-minute rest between each repetition. These training paradigms result in equal axon regeneration for each sex (25).

### Nerve collection

The animals were re-anesthetized with isoflurane. Sciatic nerves were collected at either 7- or 14-days post-injury. All animals were euthanized after nerve collection using 150 mg/kg pentobarbital sodium and phenytoin sodium (Euthasol solution from Med-Vet International prepared at 10 mg/ml in sterile saline). To measure axon profile lengths in nerves from *Thy-1-YFP-H* transgenic mice, whole nerves were fixed in 4% PFA for 30 minutes and then transferred to 0.1M PBS. Nerves from wild type mice were fixed in 4% paraformaldehyde (PFA) for 3 0 minutes and then cryoprotected overnight in 20% sucrose. For immunohistochemistry, the nerves were sectioned longitudinally using a cryostat (Leica) at 50 μm thickness. Sections were collected in 0.1M PBS pH 7.4 and processed for immunohistochemistry as free-floating sections. All sections or nerves were mounted onto slides and coverslipped using VectaShield mounting medium with or without the nuclear stain DAPI (Vector Labs), respectively. Slides were stored at 4°C until imaging.

### Immunohistochemistry

Nerve sections were blocked for 1 hour using 10% normal donkey serum (EMD Millipore, Catalog no. 566460) in 0.1M PBS pH 7.4, 0.3% Triton100. The following primary antibodies were diluted in blocking solution, and the nerve sections were incubated for 24 hours at room temperature with gentle rocking: mouse anti-TUJ1 for microtubules in the axons (1:1000, Biolegend Catalog No. 801202), rat anti-PECAM-1 for endothelial cells (1:1000, Biosciences, Catalog No. BDB553369), rabbit anti-IBA1 for macrophages (1:1000, Abcam Catalog no. ab178846), or rat anti-CD68 1:500 for macrophages (Abcam, Catalog No. ab53444). The nerve sections were then washed three times for 10 minutes with 0.1 M PBS at room temperature with light shaking. Biotin donkey anti-rat (1:200, Abcam Catalog No. ab102180) in 0.1M PBS was applied for 2 hours at room temperature in the dark with gentle rocking. The nerve sections were then washed three times for 10 minutes with 0.1M PBS. Secondary antibodies used: donkey anti-mouse TRITC (1:200, Abcam, Catalog No. ab6817), donkey anti-rabbit 488 1:200 (Immunoreagents Inc, Catalog No. 10151-700), Streptavidin 647 (1:200, Invitrogen Catalog no. S32357) in 0.1M PBS was then applied for 2 hours at room temperature in the dark with gentle rocking. The nerve sections were finally washed three times for 10 minutes in 0.1M PBS, air dried for 30 minutes in the dark and coverslipped with VectaShield containing the nuclear stain DAPI (Vector Labs).

Images of the injury site were obtained using an upright florescent microscope (Leica DM 6000B) at 20X magnification and were analyzed in FIJI. The region of interest (ROI) was defined as the injury site plus 1 mm rostral and caudal. Images were also obtained from approximately the same area in intact nerves. Macrophages that were reactive to CD68 and IBA-1 antibodies were identified by using the thresholding function in FIJI. The watershed function was applied, and cells within the ROI were counted using the count particles function with the following size and circularity parameters: IBA1 45-450, 0.5-1; CD68 80-450, 0.5-1; DAPI 5-Infinity, 0.5-1. The number of IBA1+, CD68+, or DAPI+ cells were counted in 5-10 images from each nerve. To quantify blood vessels, a threshold was established on PECAM-1 immunoreactivity, the ROI tool was used to select the nerve, and the % area of fluorescent PECAM-1 was measured. All processing was performed while comparing to the original images.

### In vivo angiogenesis inhibitor

Cabozantinib is an inhibitor of vascular endothelial growth factor receptor (primarily VEGF-R2) and inhibits angiogenesis after nerve transection by blocking endothelial cell recruitment to the tissue bridge when administered on day 3 post-injury (5). If given after vascularization has already occurred (day 5 post-injury), the vessels remain intact. Here, we asked whether exercise treatment, which robustly enhances axon regeneration, could overcome the lack of vascularization, e.g., axons might utilize the fibrin glue as a matrix to cross the tissue bridge. Wild type mice were orally administered a dose of 100 mg/kg of cabozantinib on day 3 (pre-vascularization, n=4 mice) or day 5 (post-vascularization, n=4 mice) (5) after nerve transection and fibrin glue repair. All mice were exercised beginning on day 3 as described above. Nerves were harvested at 7 days post-injury to assess vascularization and axon regeneration via antibody reactivity to mouse anti-TUJ1 for microtubules in axons (Biolegend Catalog No. 801202) and rat anti-PECAM-1 for endothelial cells within vessels (Biosciences, Catalog No. BDB553369).

### Axon length analysis

The regenerating axons from a total of 29 Thy-1-YFP-H nerves were imaged on an Olympus FV1000 confocal microscope at low magnification (10x, z step: 1.5 μm) using the EYFP (488) laser. The images were tiled and stitched to reconstruct the proximal and distal regenerating axons in three dimensions. Images were then loaded into FIJI. Using the straight-line tool with a diameter of 5 mm, a ROI was placed at the injury site. Using the freehand tool with a diameter of 1 mm, each axon profile length was measured starting from the ROI at the injury site and scrolling through the entire z stacks to its distal tip. The lengths of regenerating axons were sorted into bins of increasing lengths. The median axon length was calculated for each nerve as well as the cumulative percentage of the total number of these regenerating axons. The number of parent axons in the proximal stump (proximal to the injury site) was counted. This number was expressed as a ratio to the number of axons in the distal stump (distal to the injury site) to estimate the degree of axon branching.

### Electromyography

Surgery and/or treadmill training was performed as described above. The extent of axon regeneration and functional muscle reinnervation was investigated at 4 weeks post-injury. In isoflurane anesthetized mice (n=32; 10 untreated intrinsic, 6 untreated fibrin glue, 6 exercised intrinsic, 10 exercised fibrin glue), the injured sciatic nerve was exposed via blunt dissection and an electrical stimulating cuff was placed around the entire sciatic nerve proximal to the original injury. The bipolar stimulating electrode was assembled from a short length of Silastic tubing and stranded stainless steel microwire, AWG size 40 (Cooner Wire AS631). Fine wire electromyography (EMG) electrodes (California Fine Wire Company, MO#M468240), in which the insulation was removed from the distal 1 mm of the recording tips, were placed into the gastrocnemius muscle. Electrically evoked EMG activity was recorded from these electrodes in response to sciatic nerve stimulation (0.1 ms electrical pulses). Short latency direct muscle (M) responses (or compound muscle action potentials, CMAPs) are produced by motor axon activation. Stimulus intensity (ranging from 0.1V to 4V) was incrementally increased until a maximal amplitude direct M response was recorded. The maximal M response was assumed to represent the maximum extent of functional motor reinnervation of the muscle. Using custom LabView software (described here (34)), the average rectified voltages in a defined M response time window were quantified for each mouse. The M responses were recorded prior to injury and again four weeks following nerve transection and repair. The maximum M response amplitudes are scaled as a percentage of the M responses of intact to represent a measure of recovery.

### Muscle and neuromuscular junction histomorphometry

In addition to functional EMG, muscle reinnervation was also assessed anatomically by counting re-innervated motor endplates in gastrocnemius muscle sections from an additional 17 wild type mice. The number of presynaptic terminals immunoreactive to neurofilament 200 and synaptic vesicle 2 was assessed four-weeks following nerve transection and repair. Intact and re-innervated gastrocnemius muscles were harvested and cryoprotected in sucrose overnight. Muscles were then sectioned on a cryostat in a horizontal plane at 20 µm thickness, placed on slides, and reacted with an antibody to synaptic vesicle 2 (SV2 1:50; Developmental Studies Hybridoma Bank Ab2315387) and neurofilament 200 (NF200 1:500, Abcam Ab72996) for 24 hours at 4°C, followed by a goat anti-mouse and goat anti-chicken immunoglobulin secondary antibodies conjugated to Alexafluor 488 (1:200; Invitrogen A32931 and A32723), and rhodamine-conjugated α-bungarotoxin (1:1000; Invitrogen B35451). In each muscle studied (17 mice, 3-5 mice per group), fluorescent images of 180 motor endplates (on average for each case) were obtained and scored for immunoreactivity (IR) to SV2/NF200 within the boundaries of the endplate which was marked by bungarotoxin binding to the acetylcholine receptor (AChR). The presence of presynaptic SV2/NF200 within the endplate indicated the nerve terminal within the neuromuscular synapse. The morphology of the synapses were also analyzed using NMJmorph for the following post-synaptic morphological variables: AChR perimeter (μm), AChR area (μm^2^), endplate diameter (μm), endplate perimeter (μm), compactness (%), endplate area (μm^2^), and fragmentation. NMJmorph is an open access, ImageJ-based package for morphometric analysis of neuromuscular junctions (NMJs) and has been published with detailed methodological steps (35, 36). Care was taken to analyze only *en face* NMJs. Confocal images were obtained on a Nikon A1R HD using 512 × 512 frame size, 63X objective, 2X zoom, and 1 μm z-stack interval.

### Statistics

All results were scored while blinded to treatment and surgery repair type. The normality of residuals was analyzed for each data set using the Anderson-Darling omnibus test as well as visual inspection of the QQ plots. The average median axon lengths, branching ratios, M response amplitudes, muscle re-innervation (SV2/NF IR), blood vessel density, and CD68 cell counts were compared between groups using a one-way ANOVA with Tukey’s or Fisher’s LSD post-hoc testing. Cumulative axon lengths, IBA1 cell counts, DAPI cell counts, and the duration of the direct muscle responses were not normally distributed and thus were analyzed using the Kruskal-Wallis ANOVA with Dunn’s multiple comparisons post-hoc testing or the Mann-Whitney U according to group size. Blood vessel density after cabozantinib treatment was analyzed with an unpaired Student’s t-test. Axon length after cabozantinib treatment was compared between groups using a two-way ANOVA (mixed model) corrected for multiple comparisons. Principal components analysis (PCA) was performed for post-synaptic NMJ variables. A total of 495 NMJ profiles were incorporated into the data set from 4-week post-injury gastrocnemius muscle (intact, n=99; unexercised intrinsic repair, n=98; unexercised fibrin glue repair, n=97; exercised intrinsic repair, 112; and exercised fibrin glue repair, n=89). PCA is a data reduction method that uses structure detection to express two or more correlated variables with one or more uncorrelated variables that are termed principal components. The proportion of overall variance in a data set that is explained by each principal component is termed its eigenvalue. To obtain the maximum variance explained by each principal component, eigenvalues were obtained using the Kaiser rule. The first principal component accounts for the largest proportion of the variance in the data and has the largest eigenvalue. Each succeeding principal component explains the maximum amount of the remaining variance and is uncorrelated with previous principal components.

Principal component (factor) loading values resulting from PCA were used in subsequent statistical analysis. Factor loading values represent the correlation coefficient between the NMJ morphological profile and the regression line representing each principal component. They were used as a quantitative representation of the NMJ post-synaptic morphological profile (i.e., one loading value for each principal component for each NMJ profile).

For select data of interest, Cohen’s effect sizes (*d*) are also reported and were calculated as the difference in group means divided by the pooled standard deviation, where *d*=0.2 is considered a ‘small’ effect size; 0.5 is a ‘medium’ effect size; and 0.8 is a ‘large’ effect size. Statistics were computed using GraphPad Prism8.

## RESULTS

### Neither fibrin glue nor exercise alter macrophage infiltration of the injury site

Following nerve injury, immune cells invade the site and remain for days to months (37, 38). These cells can negatively or positively influence the regenerative process. We quantified the number of macrophages via CD68 and IBA1 IR in intact nerves and repaired nerves (Figure 1). The counts of IBA1-IR cells were not normally distributed, and thus a Mann-Whitney U test was performed. The numbers of IBA1-IR cells increased 7 days after injury and repair compared to intact (H=12.81, p < .05), but counts of IBA1-IR cells did not differ significantly between the four injury groups (Figure 1M). The number of macrophages identified as CD68-IR also increased significantly 7 days after injury and repair compared to intact (F _4,14_ = 6.326, p < .004), but counts of CD68-IR cells did not differ significantly between the four injured groups (Figure 1N). The overall number of cells present as indicated by DAPI counts also did not differ between injury groups, although nerves from all four injured groups exhibited significantly more cells compared to nerves from intact uninjured mice (data not shown). These data suggest that neither fibrin glue nor exercise treatments significantly alter numbers of macrophages 7 days post-injury.

**Figure 1.**
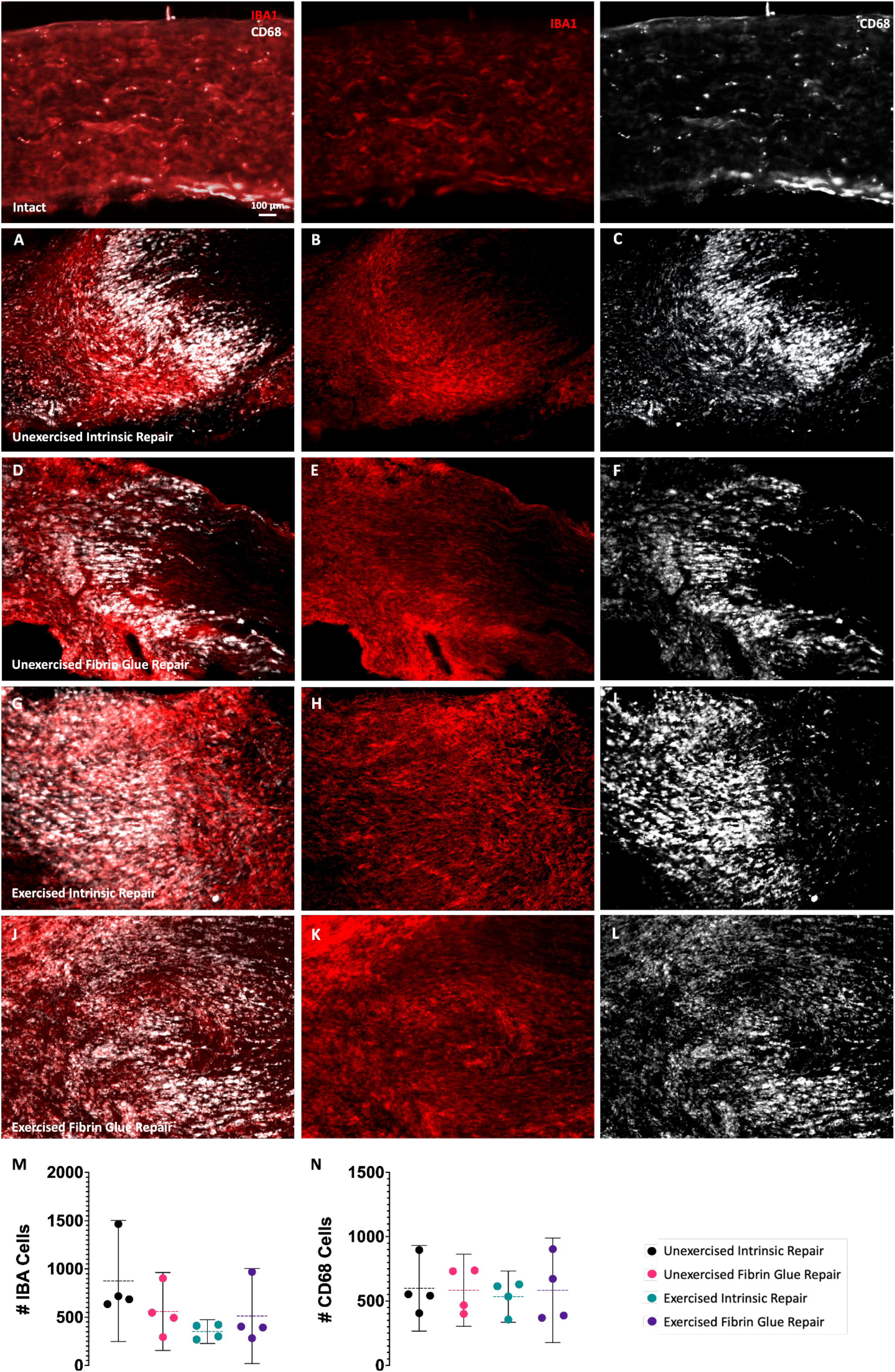
Effect of fibrin glue on macrophage infiltration. Macrophages were visualized by IBA1 (red) and CD68 (gray) immunoreactivity. At 7 days post-injury, many more cells were present within the injury site and proximal and distal nerve stumps. There were no differences in the numbers of IBA1 or CD68-IR cells among the injury groups. The means and 95% confidence intervals of IBA1 and CD68 IR cells are shown as adjusted to Intact (M-N).

### Fibrin glue and exercise increase vascularization of the injury site

It is well appreciated that neovascularization occurs after an injury and is necessary to support axon growth and prevent necrosis (39, 40). The extracellular matrix plays a large role in the development of these new vessels, and the components of fibrin glue (fibrin, thrombin, and fibronectin) can provide a matrix as well as direct endothelial cell migration through biological signaling (41, 42). An antibody against platelet endothelial cell adhesion molecule-1 (PECAM-1) was used to mark endothelial cells within the vascular compartment. To facilitate visualization of the repair site, TUJ1 (beta-tubulin) was used to identify the axons. The percent area of the nerve injury site with PECAM-1 immunoreactivity was used to quantify blood vessel density within the tissue bridge (Figure 2). At 7 days after injury, new blood vessels were observed within the tissue bridge in all injury groups, and the amount of PECAM-1 immunoreactivity was greater in the tissue bridge of injured nerves than observed within intact nerves (Figure 2E-F). The omnibus test of the ANOVA of the percent area of PECAM-1 fluorescence was significant (F _4,14_ = 27.25, p < .0001). Significantly more vessels were present in nerves repaired with fibrin glue compared to intrinsic repair (p < .01). The exercise fibrin glue group exhibited significantly more vessels than all other groups (p < .01). These data suggest that fibrin glue and exercise may work synergistically to increase acute angiogenesis (7 days post-injury).

**Figure 2.**
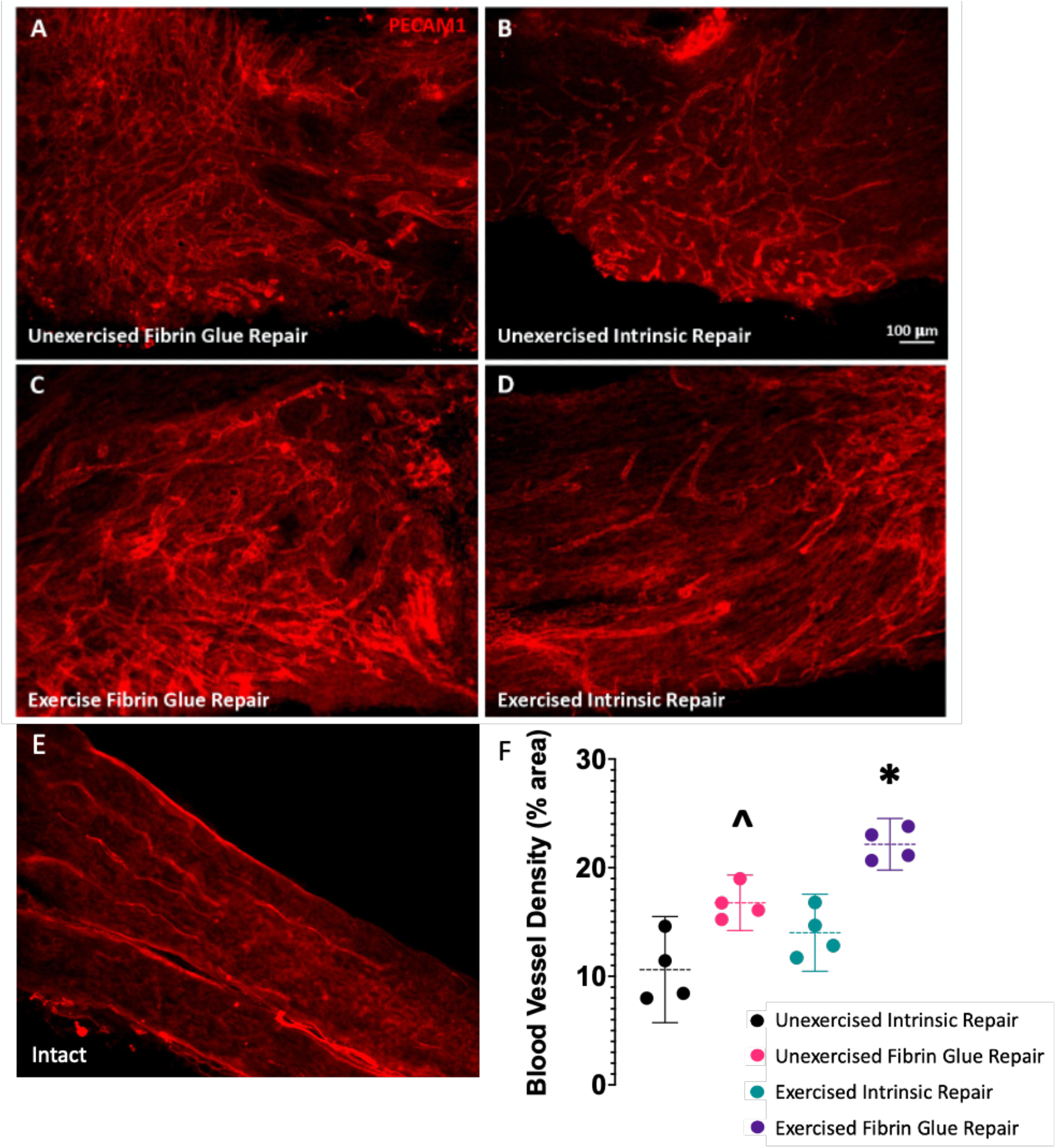
Effect of fibrin glue on vascularization. Vasculature was visualized by PECAM-1 immunoreactivity (red). At 7 days post-injury, new blood vessels were present within the injury site (A-D). Fibrin glue repaired nerves exhibited significantly more vessels than the intrinsic repair. The combination of exercise and fibrin glue repair resulted in the greatest increase of vasculature in the tissue bridge (C). The means and 95% confidence intervals blood vessel density is shown as adjusted to Intact (F). *Exercise Fibrin Glue Repair vs all groups. ^Unexercised fibrin glue vs unexercised intrinsic repair.

### Angiogenesis is critical for the regenerative effects of exercise

Next, we tested our hypothesis that increased vascularization of the fibrin glue tissue bridge was required for the enhanced axon regeneration in exercising mice. We used an inhibitor of endothelial cell recruitment and tube formation to disrupt angiogenesis in the tissue bridge. Angiogenesis of the nerve bridge occurs between days 3 and 5 after nerve injury, and cabozantinib treatment on day 3 (pre-vascularization) blocks vascularization of the bridge and subsequently prevents axon crossing (5). In exercising mice with fibrin glue nerve repair, we found a lack of vascularization of the nerve bridge when mice were treated with cabozantinib on day 3 (pre-versus post-vascularization, t(6) = 3.263, p < .05) and a reduction of axon growth into the distal nerve stump (Figure 3A and C) (ANOVA, F _1.28,9.43_ = 56.15, p < .0001). In contrast, if exercising mice were treated with cabozantinib on day 5 (post-vascularization), vascularization of the bridge and axon regeneration proceeded (Figure 3B and C). Although two weeks of exercise is known to robustly enhance peripheral axon regeneration through multiple mechanisms, vascularization of the tissue bridge is required for the enhancing effect of exercise (Figure 3D).

**Figure 3.**
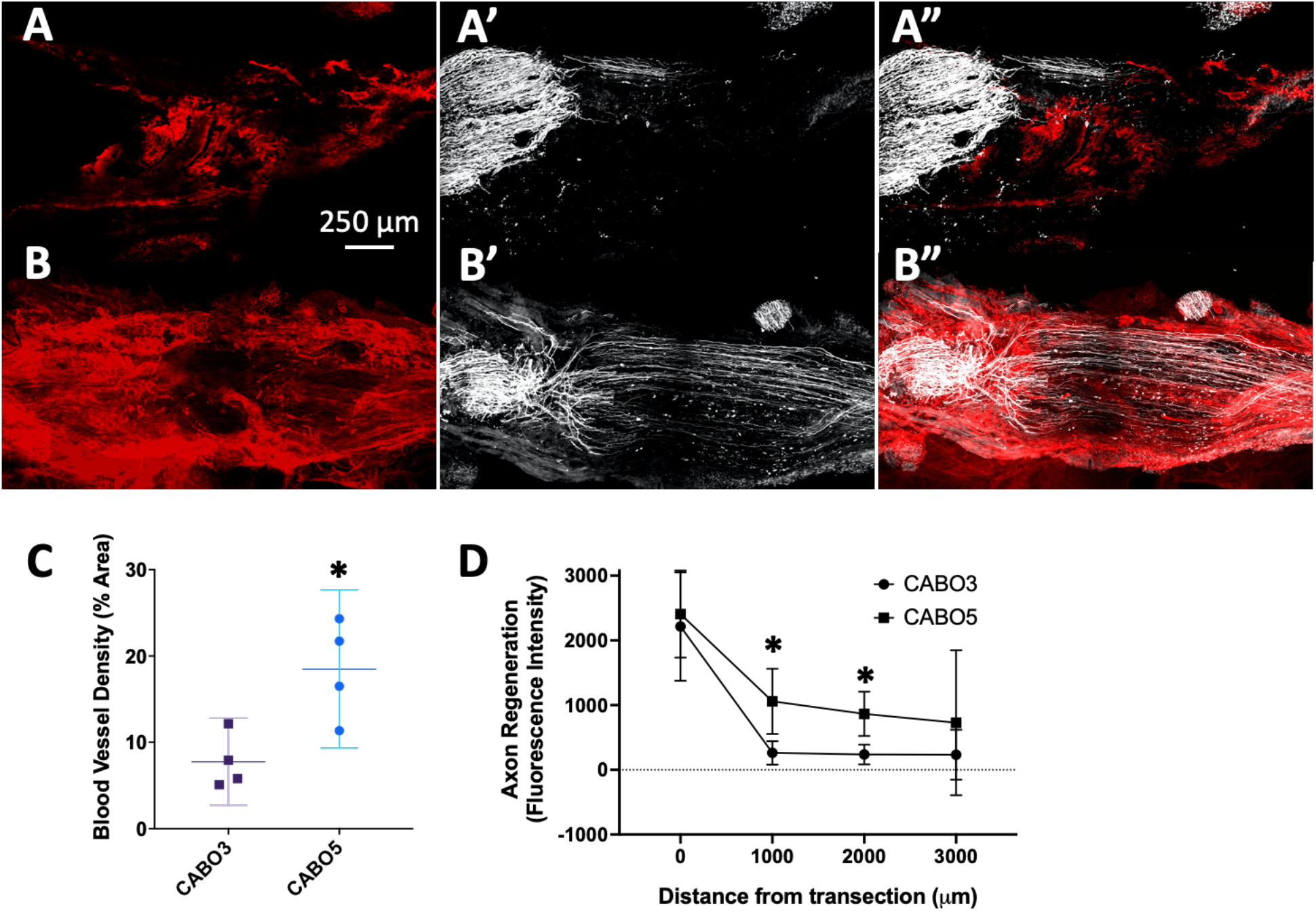
Effect of fibrin glue and exercise on axon regeneration in the absence of angiogenesis. Representative images of the nerve bridge 7 days post-injury. Axons and vasculature were visualized with TUJ1 and PECAM-1, respectively. When cabozantinib (an inhibitor of vascular endothelial growth factor receptor) was administered on day 3 post-injury, angiogenesis was inhibited after nerve transection and few axons crossed the injury site despite the combination of fibrin glue repair and exercise treatment (A-A’’). If cabozantinib was administered on day 5 post-injury (after vessels have formed), many axons cross the injury site (B-B’’). C) Quantification of blood vessel density. D) Quantification of axon growth into the distal stump. Means and 95%CI are shown (C-D).

### Fibrin glue repair combined with exercise results in the greatest axon elongation

Angiogenesis after injury is a necessary step that occurs prior to axon elongation (5, 43). Given our observation of greater vasculature within exercised fibrin glue nerves (Figure 2) and that axon elongation was prevented by inhibiting angiogenesis (Figure 3), we next asked whether axon elongation was significantly improved with exercise in nerves repaired with fibrin glue using our well characterized transgenic model of regeneration (25, 27, 32, 33, 44). To visualize the growing axons, we used *Thy-1-YFP-H* mice that express YFP in a subset of axons (Figure 4). Lengths of individual axon profiles of fluorescent regenerating axons were measured using optical sections made through whole mounts of harvested nerves. We examined the median axon length of each nerve as a measure of central tendency. Significant differences in median axon profile lengths were noted (ANOVA, F _3,25_ = 3.499, p < .05). In mice whose nerves were repaired with fibrin glue and exercised, the average median axon profiles were significantly longer than similarly repaired untreated mice (Cohen’s *d* = 1.07) and untreated intrinsic repair mice (Cohen’s *d* = 1.31) (LSD, both p = .01) (Figure 4E). Interestingly, axon branching was significantly greater in the untreated intrinsic repair group (Figure 4F), which may indicate a lack of guidance cues. Lastly, the cumulative distributions of axon profile lengths were not normally distributed; thus, we performed Kruskal-Wallis ANOVA to test whether the samples originated from the same distributions. The distribution of axon lengths from exercised mice with fibrin glue repair were significantly shifted to the right ((χ^2^ = 12.63, p < .01) compared to untreated mice with and without fibrin glue as well as exercised mice with intrinsic repair (all p < .05), indicating that longer axons were found in exercised mice whose nerves were repaired with fibrin glue (Figure 4G).

**Figure 4.**
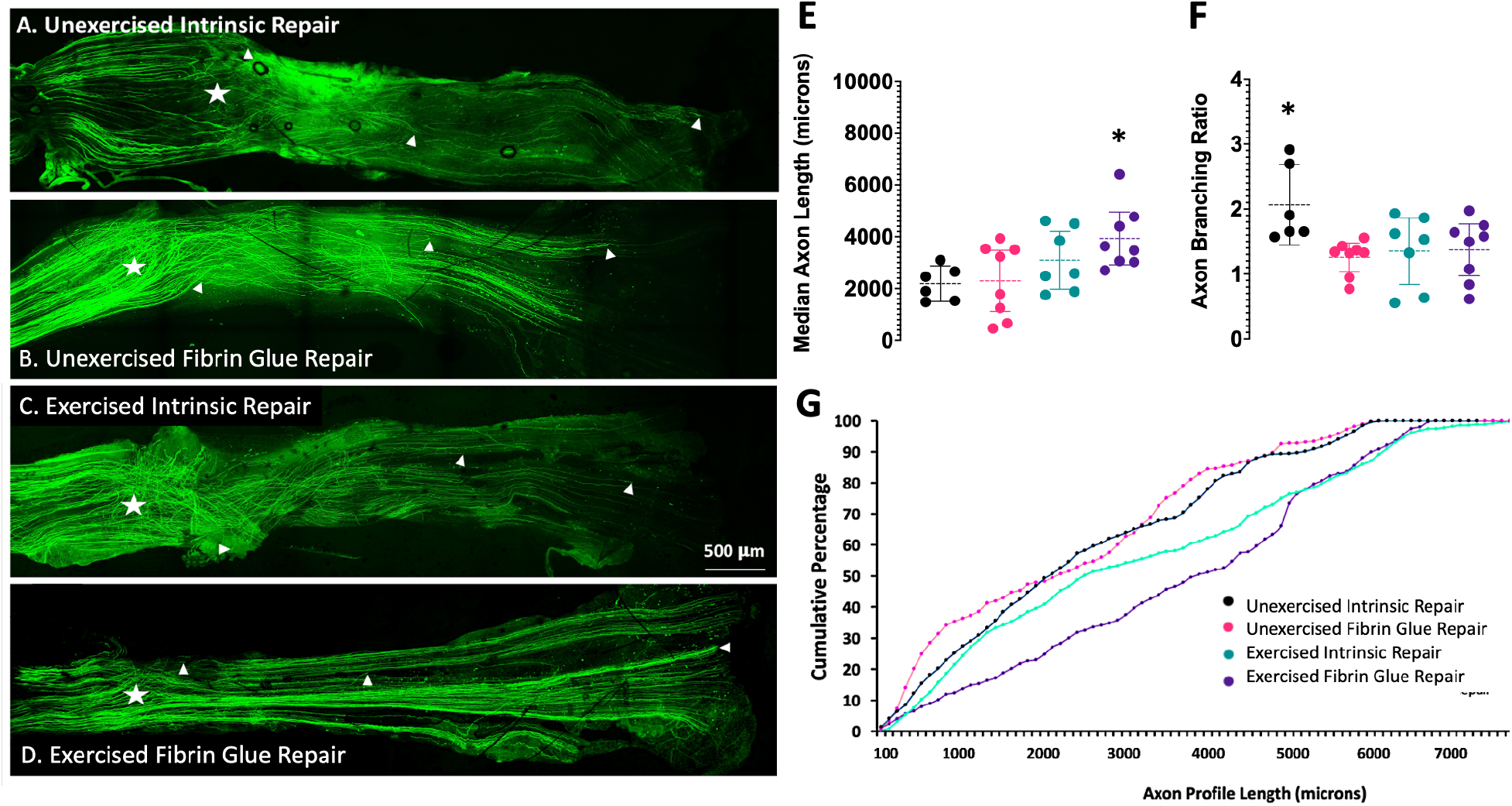
Effect of fibrin glue and exercise on axon elongation. YFP+ regenerating axons can be visualized growing into the distal stump at 14 days post-injury, and the arrowheads indicate growth cones/axon tips in A-D. The original injury site is indicated by the star. The number and length of each axon profile was recorded. E) Average (±95%CI) median lengths of regenerating axon profiles two weeks after sciatic nerve transection and repair in mice. F) The average (±95%CI) axon branching. G) The cumulative frequency distributions of each group. The cumulative distribution of axon lengths in the exercised fibrin glue repaired group are significantly shifted to the right, which indicates longer axons.

Because gap length is a well-known determinant of the success or failure of axon regeneration, we wanted to ensure that both the intrinsic and fibrin glue repair methods resulted in similar repair quality. Therefore, we measured the distance between the proximal and distal nerve stumps, e.g., the length of the nerve bridge, in the *Thy-1-YFP-H* nerves. No significant difference in gap distances was found between the four groups (untreated intrinsic repair, exercise intrinsic repair, untreated fibrin glue repair, exercise fibrin glue repair) by ANOVA (F _3,18_ = 2.304, ns). Because exercise was not expected to alter the length of the nerve bridge, we also tested for differences in gap distance by combining the untreated intrinsic repair group with the exercise intrinsic repair group, and the untreated fibrin glue repair group was combined with the exercise fibrin glue repair group. These groups were tested for significant difference using a two-tailed independent t-test, and no significant difference in gap size was noted, *t*(20)=1.007, ns. Both methods of nerve injury and repair resulted in similar gap lengths (mean intrinsic repair mean = 752.2 μm 95% CI [655-848]; fibrin glue repair mean gap length = 885.9 μm 95% CI [507.7-1264].

### Exercise with fibrin glue results in greater functional recovery and neuromuscular re-innervation

To evaluate whether the acute effect of fibrin glue on enhancing axon elongation resulted in greater functional recovery, compound muscle action potentials (CMAPs or M responses) were recorded from gastrocnemius muscle in response to proximal sciatic nerve stimulation on same mice prior to and four weeks after sciatic nerve injury (Figure 5A). Amplitudes of the largest electrically evoked M responses are displayed as a percentage of intact values that were recorded prior to nerve transection. The evoked motor response amplitudes were significantly greater in exercised mice with fibrin glue repaired nerves compared to all other groups (ANOVA, F _3,28_ = 13.14, p < .001). In exercised mice whose nerves were repaired with fibrin glue, the mean evoked EMG amplitude was restored to ∼32% of intact (Figure 5B). The M responses from exercised mice whose nerves were not repaired with fibrin glue were more variable, and the mean evoked EMG amplitude was restored to ∼18%. The M responses from mice in both untreated groups (fibrin glue or without) indicated little functional motor recovery, reaching 10% or less (Figure 5B). M response amplitudes from the exercise fibrin repair groups were significantly greater than untreated fibrin repair and untreated intrinsic repair groups (p < .001) as well as exercise intrinsic repair (p < .05). Thus, compound muscle action potentials are restored to a greater extent in exercised mice whose nerves were repaired with fibrin glue compared to all others (Cohen’s *d* = .78). The duration of the M response was decreased in the exercised fibrin glue repair group (Figure 5D), which may represent compound muscle action potential maturation (45).

**Figure 5.**
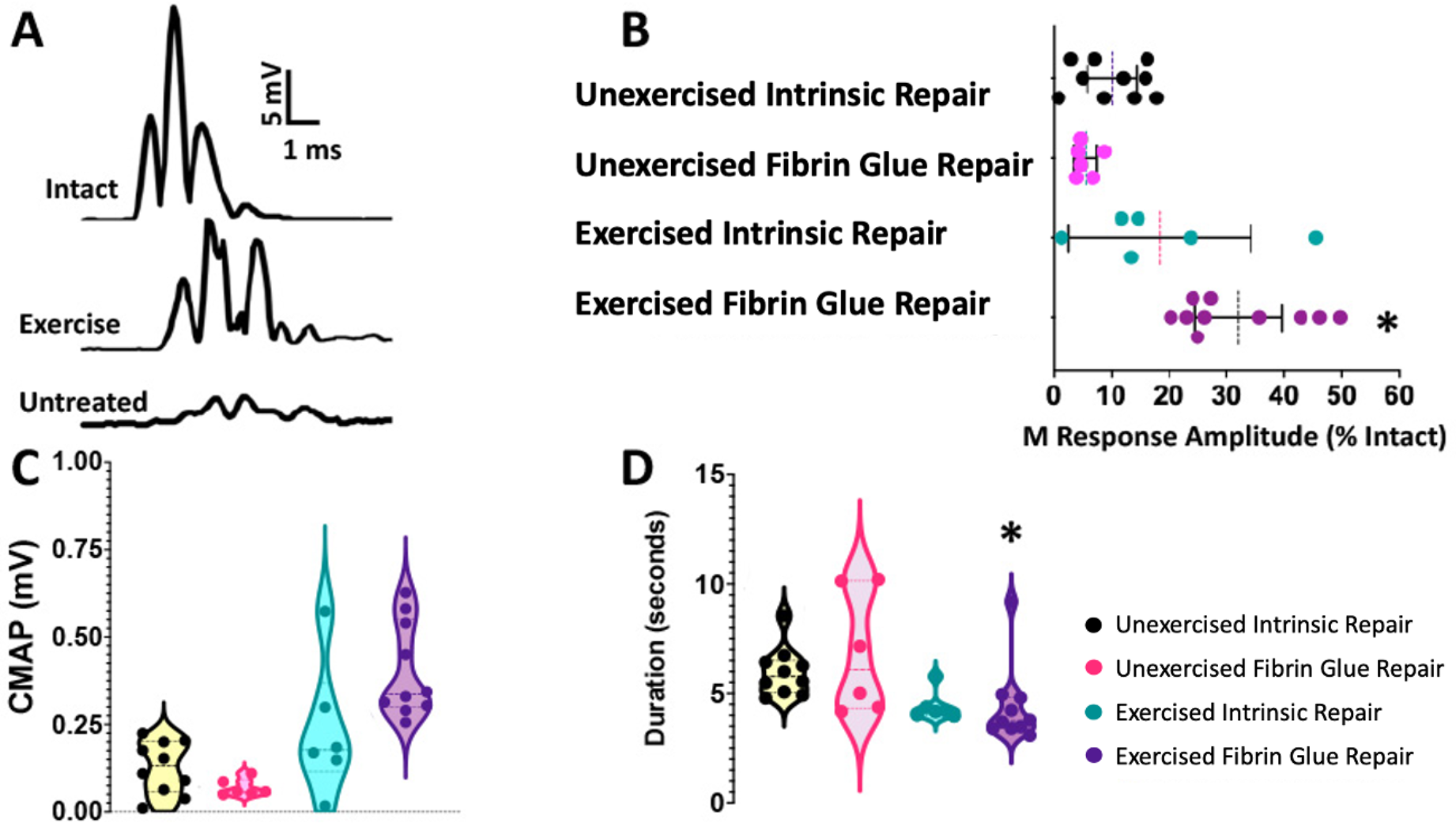
Effect of fibrin glue and exercise on functional recovery. Representative traces of evoked EMG activity in the gastrocnemius muscle. Electrically-evoked direct muscle responses represent functional motor axons that have successfully reinnervated the muscle. A) Typical intact M response vs exercised and untreated mice whose nerves were repaired with fibrin glue. B) Means and 95%CI of M response amplitudes (4 weeks post-injury) scaled as a percent of the mean intact amplitude. *Exercise Fibrin Glue Repair vs all groups. C) Compound muscle action potentials in absolute values. D) Duration of direct muscle response. The duration shortens as the M response matures during re-innervation. * Exercise Fibrin Glue vs Untreated Intrinsic Repair.

In a separate group of mice, the extent of muscle re-innervation was assessed by confocal microscopy of NMJs at four weeks after injury. The presence or absence of a nerve terminal within each motor endplate was assessed as indicated by presynaptic SV2/NF (Figure 6A and 6B). By four weeks post-injury and repair (ANOVA, F _4,12_ = 97.92, p < .001), the exercised fibrin glue group exhibited significantly greater numbers of re-innervated NMJs compared to all other treatment groups, although still less than intact levels (all post-hoc p < .001, Figure 6C). Next, we used factor analysis with principal components (PCA) to evaluate the significance of any changes in post-synaptic morphology following gastrocnemius muscle reinnervation between treatment groups at four weeks (Figure 7A-E). The scree plot is shown in Figure 7F. The first two principal components explained 81.26% of the variance in the data set (Figure 7G). The first principal component PC1 (primarily related to the “overall size” of the NMJs) alone explained 65.48% of the variance. Both PC1 and PC2 (primarily related to the compactness of NMJs, as indicated by the positive correlation and position on the y coordinate on the PCA loading plot) had eigenvalues that were greater than 1.0, which means that each of them alone explained a much larger proportion of overall variance than any single raw variable. The eigenvalue for PC3 fell below 1 and explained only 9.08% of the variance in the data set. Therefore, we only considered PC1 and PC2 in further analysis. Factor loadings, the correlation of eigenvalues to the original data set, were calculated for each PC for each group. The significance of the differences in factor loadings between the different groups was determined using ANOVA and post hoc testing.

**Figure 6.**
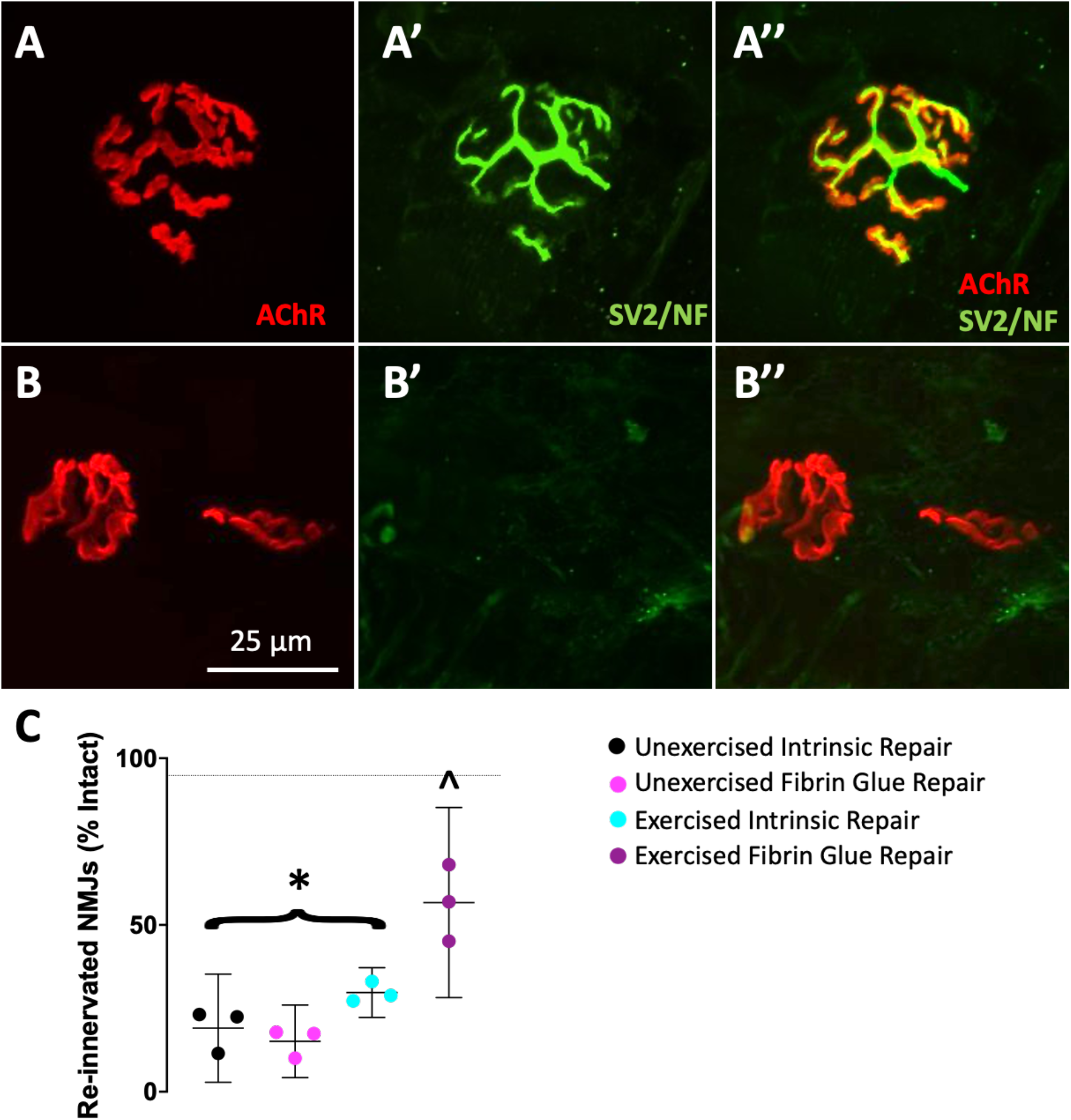
Effect of fibrin glue and exercise on neuromuscular reinnervation. Quantification of re-innervation of neuromuscular junctions. A-B) Images of neuromuscular junctions demonstrating the presence (A, A’, A”) of the axon within a reinnervated junction or absence (B, B’, B”) of a nerve terminal in a denervated terminal as indicated by presynaptic markers synaptic vesicle 2 and neurofilament 200 (SV/NF, green) within each motor endplate labeled with acetylcholine receptor (AChR, red). (C) The means and 95%CI of re-innervated NMJs are shown at four weeks post-injury and repair. The dashed line indicates the average intact value.

**Figure 7.**
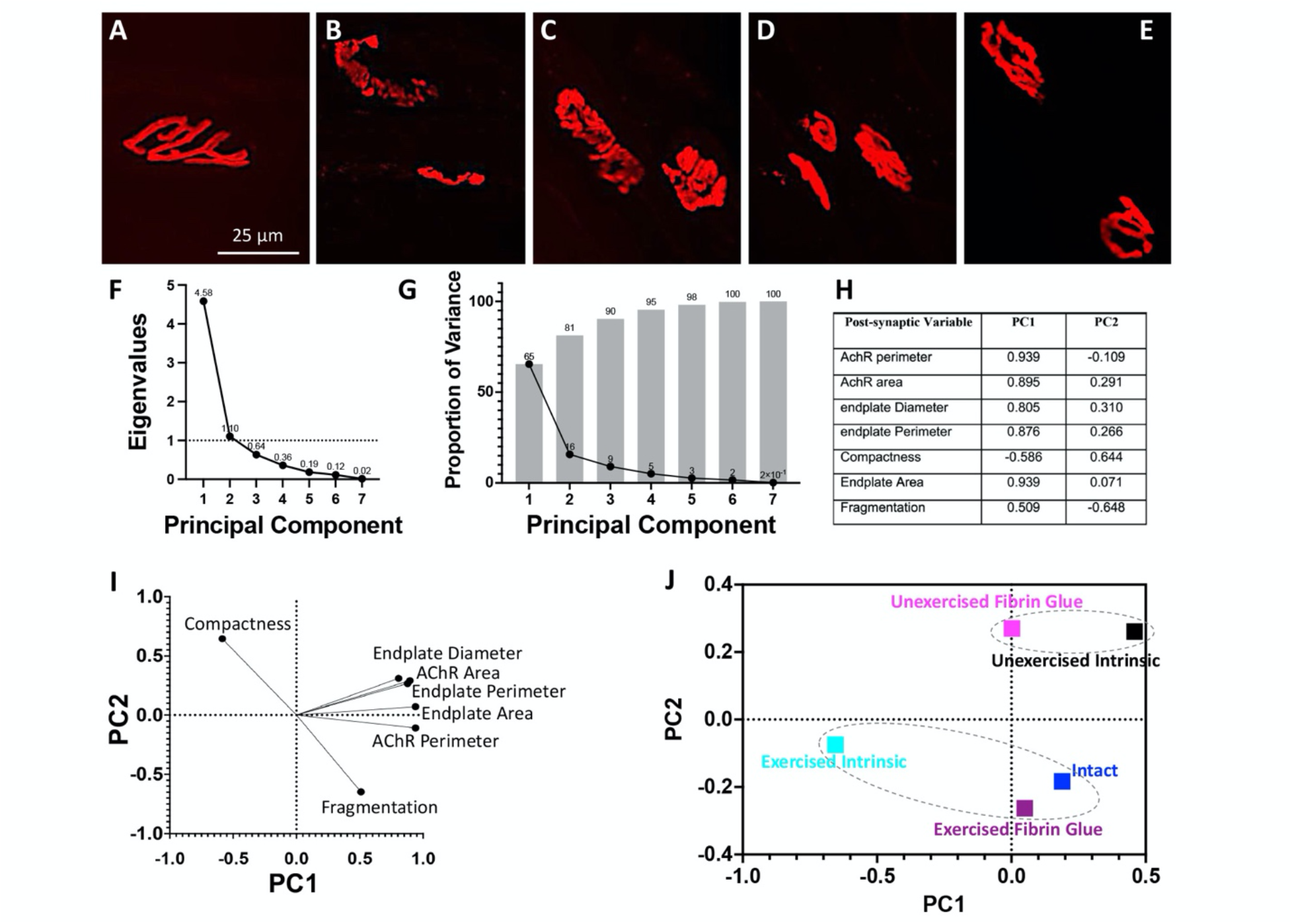
Effect of fibrin glue and exercise on neuromuscular reinnervation. Acetylcholine receptors (AChR2) within the motor endplates of neuromuscular junctions. Representative endplates from each group A) intact, B) unexercised intrinsic repair, C) unexercised fibrin glue, D) exercised intrinsic repair, E) exercised fibrin glue repair at 4 weeks post-injury. F) The scree plot from principal component analysis. G) Proportion of variance for each principal component. H) Correlation of each post-synaptic variable with PC1 and PC2. I) Relationship of each post-synaptic variable with PC1 and PC2. J) Mean loading factor scores for PC1 and PC2. Dotted lines enclose groups that are not significantly different from one another.

PC1 interpretation is based on the markedly positive correlations between the size-related morphological variables (Figure 7H) and their clustering with high x coordinate values on the PCA loading plot (Figure 7I). We next performed a Kruskal-Wallis ANOVA because the scores for PC1 and PC2 were not normally distributed. The PC1 and PC2 scores indicated significantly different distributions among groups (χ^2^ = 12.38, p < .05 and (χ^2^ = 27.13, p < .0001, respectively). The PC1 scores for the exercise intrinsic repair group were significantly different compared to untreated intrinsic repair (p < .01). While the PC2 scores for the exercise fibrin glue repair group were significantly different compared to all untreated groups (all p < .05) but were not different from intact mice. Figure 7J demonstrates these relationships by plotting the average loading factor values for PC1 and PC2 for each group. The overall conclusion we draw from the post-synaptic PCA of morphology is that at four weeks post-injury, the compactness and overall size of the motor endplate was altered by injury and recovery, and endplates observed from groups with greater re-innervation and functional recovery, e.g., exercise fibrin glue, exhibited characteristics similar to endplates found in intact mice (Figure 7J).

## DISCUSSION

Today’s surgeons repair severed peripheral nerves with needle and suture, very similar to the way it was performed a century ago (9, 46). Some surgeons prefer to use microsutures alone, some use glue or sealant alone, and some use both. Much of this choice depends on surgeon preference, the size of the nerve and characteristics of the injury. Glue alone is used more frequently in children due to small nerve calibers and difficult access, which makes microsuturing more difficult. Microsutures alone are more commonly used in adults and is considered the gold standard repair method. The majority of studies indicate no clear benefit of either microsuture versus glue, although glue may result in a faster surgery time. In our current study, we also found no enhancement of regeneration or recovery when comparing nerves repaired with fibrin glue vs without. Importantly, we observed enhanced regeneration when exercise treatment, in the form of treadmill running, which was applied after repair with fibrin glue and a lesser enhancement by exercise in nerves with a natural intrinsic repair despite similar injury gap lengths.

To our knowledge, this is the first observation of a cellular basis for fibrin glue enhancing the efficacy of an activity-based therapy (treadmill exercise) after nerve injury. The application of fibrin glue lead to greater vascularization of the nerve repair site. Exercise for as little as ten days over a two-week period, increased the elongation of axons and improved functional recovery but only if the nerve had been repaired with fibrin glue. Chen et al. observed that Bands of Büngner in an unglued nerve bridge are not wide enough to guide all of the regenerating axons (8). Cattin et al. demonstrated that vascularization of the nerve bridge (gap) is necessary for spontaneous axon regeneration and by altering the direction of the vessels, the course of axons was similarly altered (5). Although this remains to be tested, based on our observations, it seems likely that the increased vasculature in the fibrin glue repaired nerves allowed for the development of more Bands of Büngner and subsequent guidance of the exercise-enhanced axon growth. Importantly, increased vasculature alone was not enough to allow for significantly greater regeneration as we did not detect any improvement in the fibrin glue repaired group versus the intrinsic repair according to axon lengths or functional recovery. This is in agreement with multiple studies in which addition of glue was equivalent to other types of repair (47). However, when combined with exercise, the addition of fibrin glue to the nerve bridge results in greater vascularization, axon elongation, and functional recovery. These beneficial effects of fibrin glue were negated by preventing vascularization of the nerve bridge. We conclude that fibrin glue provides a microenvironment conducive to vascularization, and when combined with exercise, the increased vascularization facilitates enhanced regeneration.

There are several possible mechanisms for the increased angiogenesis found in nerves repaired with fibrin glue. Endothelial cells require extracellular matrix to migrate, and the presence of fibrin glue within the tissue bridge may act as a scaffold. Fibrin matrices are commonly used to study angiogenesis. Endothelial cells cultured on extracellular matrix with fibrin quickly form capillary-like structures but do not when cultured on plastic (48). In addition to mechanical properties, both fibrin and thrombin signal to endothelial cells through integrins, which leads to second messenger signaling and angiogenesis (23, 49). In Muhammad et al., greater numbers of endothelial cells were observed after application of a fibrin glue clot following brain injury (50). In our study of fibrin glue repaired nerves, we observed increased vasculature formation within the tissue bridge (Figure 2).

Our initial hypothesis was that fibrin glue would cause an adverse effect on regeneration perhaps due to its xenogenic nature, potential immunogenicity, and the strong chemotactic effect of thrombin on macrophages (51–53). Yet, we found no evidence to support this. There were no detectable differences in the numbers of CD68+ or IBA1+ cells within nerves repaired with or without fibrin glue regardless of exercise treatment (at 7 days post-injury), although we did not sample the inflammatory milieu. While the overall numbers of macrophages did not differ between the four treatment groups, we hypothesize that the inflammatory milieu will be affected by both fibrin glue and exercise. To confirm this hypothesis, the phenotype of the macrophages would need a careful analysis via cell sorting or other more sensitive techniques. Within the tissue bridge of nerves repaired with fibrin glue, we observed significantly more blood vessels at 7 days post-injury followed by longer regenerating axon profiles at 14 days post-injury and enhanced functional recovery at 4 weeks. Thus, one could surmise that the increased vasculature within fibrin glue repaired nerves increased the efficacy of exercise as a treatment. The efficacy of exercise after nerve surgery remains to be determined in human patients because postsurgical treatment normally restricts movement to avoid dehiscence of the surgical repair. However, brief electrical stimulation of the nerve at the time of surgery and repair can accelerate nerve regeneration in humans (54). The efficacy of electrical stimulation is well-established in preclinical models of both direct repair and long nerve grafts, and it utilizes many of the same mechanisms as exercise (24, 33, 55–59). Zuo et al. showed that a single brief session of electrical stimulation promotes both sensory and motor regeneration through long grafts placed with microsutures (60), and electrical stimulation is in clinical trials for nerve injury in humans. It would be of clinical interest to test whether electrical stimulation can further enhance nerve regeneration when combined with fibrin glue (alone or in combination with sutures), similar to our observations with exercise.

Several challenges prevent full regeneration and recovery clinically. Over the decades, the cellular orchestration of nerve repair has become clearer – involving macrophage infiltration followed by angiogenesis, Schwann cell migration along the vessels, axon elongation along the Bands of Büngner, and finally functional reinnervation of appropriate distal targets. Encouraging or mimicking a polarized vasculature within nerve grafts and conduits has been the subject of many studies. However, we may need to also consider its importance in direct nerve repairs. Even with the simplest end-to-end direct tensionless anastomosis, a microscopic gap will remain between the proximal and distal nerve stumps, and it has long been appreciated that nerve gaps represent a major challenge for nerve regeneration, and functional recovery can take years after nerve repair (46, 61). Long nerve gaps are often repaired with nerve grafts or conduits, and little regeneration occurs compared to direct tensionless anastomosis. In part, this poor regeneration is due to the need to cross two gaps – proximal nerve stump to graft then graft to distal nerve stump - and the addition of fibrin glue at each gap may support the crossing of more axons. In our results, even a microscopic gap (∼800 μm) represented a major challenge to the regenerating axons. The addition of fibrin glue in this gap supported greater angiogenesis which then served as an adjuvant for exercise, an activity-based therapy, resulting in greater functional recovery. Although many tissue glues and sealants are now available (for reviews see (12, 62)), the biological properties of fibrin glue may be superior in nerve repair not for its mechanical stability (which is weak) but for its ability to support angiogenesis, which in turn supports enhanced growth when combined with a treatment that enhances axon regeneration.

## Conclusion

Numerous studies have compared fibrin glues versus sutures with no definitive benefit, leaving nerve surgeons to rely primarily on microsutures due to the risk of dehiscence with glue alone unless necessitated by small nerve caliber or lack of accessibility (10, 11). In this study, angiogenesis was critical for the regenerative effects of exercise, and fibrin glue supported greater vascularization within the tissue bridge resulting in enhanced functional efficacy of treadmill exercise. Our results suggest that fibrin glue may positively support intra- or postoperative physical rehabilitation or activity-based therapies that enhance axon regeneration via increased vascularization of the tissue bridge. The addition of fibrin glue – even if microsutures are the primary repair strategy – is therefore a promising and clinically readily available adjuvant to rehabilitation strategies for a patient population which currently has no effective, non-surgical treatment options. Therapies that encourage early vascularization of the gap after transection should be considered even when a tensionless direct repair can be achieved.

## Acknowledgments

We would like to thank Dr. Arthur English for his constructive comments on the manuscript, and Dr. Andrew Kowalczyk for providing the PECAM-1 antibody. This research project was supported by the Emory University Integrated Cellular Imaging Core; the National Center for Advancing Translational Sciences of the National Institutes of Health under Award Number UL1TR002378; the NIH Eunice Kennedy Shriver National Institute of Child Health and Human Development, National Center for Medical Rehabilitation Research under award number HD09770737, and in part by a developmental grant from the NIH-funded Emory Specialized Center of Research Excellence in Sex Differences U54AG062334 to PJW.

## Author Contributions

PJW contributed to the conception and design of the study; SSW, ADB, TT, TSP and PJW contributed to the acquisition and analysis of data; SSW, ADB, and PJW contributed to drafting the text and preparing the figures. All authors revised and approved the final manuscript.

